# Gap selection and steering during obstacle avoidance in pigeons

**DOI:** 10.1101/2022.03.01.482520

**Authors:** Natalia Pérez-Campanero Antolin, Graham K. Taylor

## Abstract

The ability of birds to fly through cluttered environments has inspired biologists interested in understanding its underlying mechanisms, and engineers interested in applying its underpinning principles. To analyse this problem empirically, we break it down into two distinct, but related, questions: How do birds select which gaps to aim for? And how do they steer through them? We answered these questions by using a combined experimental and modelling approach, in which we released pigeons *(Columbia livia domestica)* inside a large hall whose open exit was separated from the release point by a curtain creating two vertical gaps—one of which was obstructed by an obstacle. We tracked the birds using a high-speed motion capture system, and found that their gap choice seemed to be biased by their intrinsic handedness, rather than determined by extrinsic cues such as the size of the gap or its alignment with the destination. We modelled the pigeons’ steering behaviour algorithmically by simulating their flight trajectories under a set of six candidate guidance laws, including those used previously to model targeted flight behaviours in birds. We found that their flights were best modelled by delayed proportional navigation commanding turning in proportion to the angular rate of the line-of-sight from the pigeon to the midpoint of the gap. Our results are consistent with this being a two-phase behaviour, in which the pigeon first heads forward from the release point before steering towards the midpoint of whichever gap it chooses to aim for under closed-loop guidance. Our findings have implications for the sensorimotor mechanisms that underlie clutter negotiation in birds, uniting this with other kinds of target-oriented behaviours including aerial pursuit.

## INTRODUCTION

When B.F. Skinner proposed using pigeons to guide flying vehicles in World War II (9), he may have been onto something. Pigeons have colonized complex, cluttered urban environments throughout the world, which they negotiate successfully at high speeds. They achieve this visually, aided by their panoramic (300*°*) field of view and visual processing some three times faster than a human’s (17). However, rather than using operant conditioning to train pigeons to pilot vehicles by pecking at a screen as Skinner proposed, a better approach might have been to study the guidance algorithms by which they steer their flight. Here, we set out to do just that, using a combined experimental and modelling approach to investigate how pigeons steer towards gaps. This work is closely inspired by previous research on pigeons flying through a dense forest of vertical poles (20), but aims to reduce the problem to its simplest level, by presenting the birds with a binary choice between an obstructed or unobstructed gap through which to fly.

Algorithmic approaches to the study of animal behaviour (16) have been successful in explaining the detailed flight trajectories of bats (13), raptors (6; 8; 7), and flies (11) when intercepting prey, and pigeons when negotiating clutter (20). These studies use differential equations to build simple phenomenological models capable of accurately describing complex behavioural data. Such models exploit the goal-directed nature of target-oriented flight to identify the most relevant input variables from among the small set of candidates that can possibly be used to command turning. Typically, the angular direction or angular rate of the line-of-sight from the subject to its target is measured with respect to either its velocity vector or an inertial reference (Fig. 1), and used to command steering. Each such input variable is assumed to be fed back in proportion to its own guidance gain, which is what determines its effect upon steering. Identifying which input variables best model an animal’s steering output therefore has important implications for understanding the underlying physiological mechanisms. The resulting input-output relationship is known as a guidance law (see Table 1). Importantly, because it is only the motion of the subject relative to its target that matters in target-oriented flight, the same guidance laws can be applied to either stationary or moving targets. This opens the possibility of unifying target chasing and clutter negotiation behaviours under one common algorithmic framework.

**Table 1.** Guidance laws used previously or here to model target-oriented steering behaviour in animal flight. See text for definitions of terms.

| Guidance law | Inputs | Name | Examples |
| --- | --- | --- | --- |
| $\dot{\gamma}(t) = k_N \dot{\lambda}(t - \tau)$ | 1 | proportional navigation | falcons (6; 7), bats (13), killerflies, robberflies (11) |
| $\dot{\gamma}(t) = k_P \delta(t - \tau)$ | 1 | pure/proportional pursuit | blowflies (5), tiger beetles (15), honey bees (33) |
| $\dot{\gamma}(t) = k_P [\delta(t - \tau) - \delta_0]$ | 1 | deviated pursuit | hoverflies (12), bluefish (21), dragonflies (23) |
| $\dot{\gamma}(t) = k_D \dot{\delta}(t - \tau)$ | 1 | derivative control | not modelled previously |
| $\dot{\gamma}(t) = k_N \dot{\lambda}(t - \tau) + k_P \delta(t - \tau)$ | 2 | mixed guidance | hawks (8) |
| $\dot{\gamma}(t) = k_P \delta(t - \tau) + k_D \dot{\delta}(t - \tau)$ | 2 | proportional-derivative control | houseflies (19) |
| $\dot{\gamma}(t) = k_N \dot{\lambda}(t - \tau) + k_D \dot{\delta}(t - \tau)$ | 2 | proportional navigation with derivative control | not modelled previously |
| $\dot{\gamma}(t) = k_P \delta(t - \tau) + k_D \dot{\delta}(t - \tau) + k_S \dot{\gamma}(t - \tau)$ | 3 | proportional-derivative control with inertia | pigeons (20) |

**Fig. 1.**
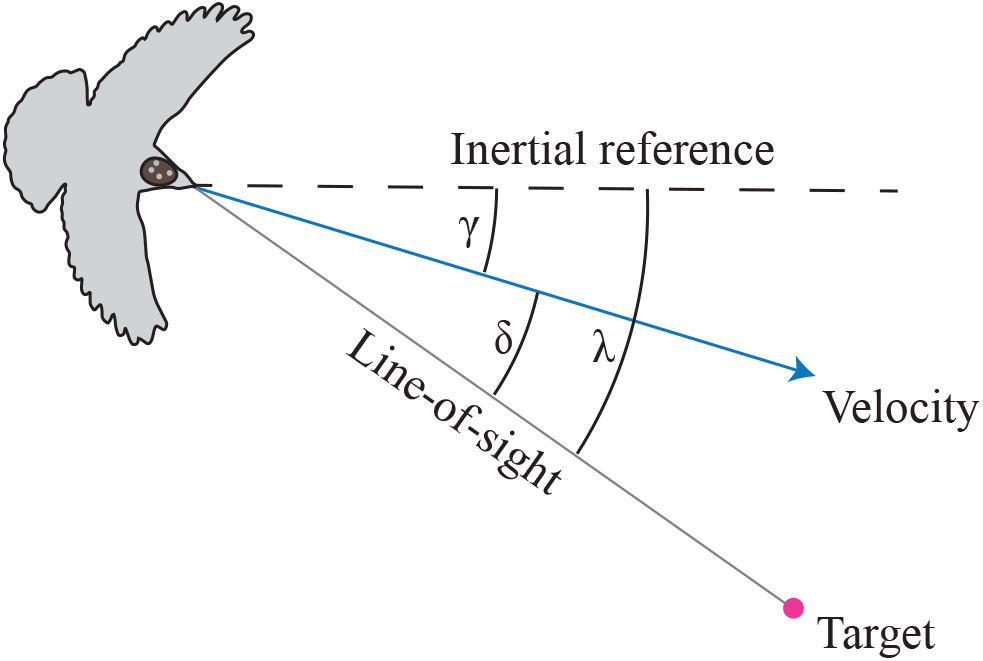
Geometry of target-oriented guidance behaviour. Definition sketch showing: the line-of-sight angle *λ* measured between the line-of-sight (grey line) from pigeon to target, and some arbitrary inertial reference direction (dashed line); the track angle *γ* measured between the bird’s ground velocity vector (blue arrow) and the inertial reference direction (dashed line); the deviation angle *δ* measured between the line-of-sight from bird to target (grey line), and the pigeon’s velocity vector (blue arrow). The arrangement of the tracked headpack markers is shown schematically.

Early attempts to model the steering of animal flight (10) looked for a proportional relationship between the subject’s turn rate 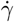 and its deviation angle (*δ*), defined as the angle between the subject’s velocity vector and its line-of-sight to the target (Fig. 1). In its simplest form, this proportionality corresponds to the guidance law 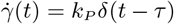, called pure or proportional pursuit (8), where *k*_*P*_ *<* 0 is the guidance gain, *t* is time, and *τ* ≥ 0 denotes a fixed time delay. Pure pursuit drives the deviation angle towards zero, thereby causing the subject to aim its flight directly at its target, but a simple variant called deviated pursuit drives the deviation angle *δ* towards some nonzero constant *δ*_0_ with 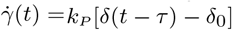. This causes flight to be aimed at a point ahead of the target, which can be effective in promoting interception over a tail chase. Since the performance of a proportional controller can often be improved by adding derivative feedback, several studies have also looked for the involvement of an additional derivative term 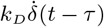, where 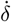 is the rate of change of the deviation angle, and *k*_*D*_ *<* 0 is the associated guidance gain (10; 20). The addition of this derivative term anticipates the changes in deviation angle that are the basis of proportional pursuit, but commands high turning rates if the deviation angle changes rapidly, which can cause instability at high gain. Because derivative control has no inherent tendency to correct for any constant offset in the deviation angle, it is usually only used in combination with proportional control, but we also test it in isolation here, to aid in statistical inference.

Other plausible steering controllers command turning in proportion to the angular rate of the subject’s line-of-sight to its target 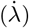, measured in an inertial frame of reference. The simplest form of this proportionality corresponds to the classical guidance law of homing missiles, 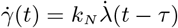, called proportional navigation (6). This single-input guidance law has been used to explain the attack trajectories of falcons (6; 7), and effectively combines a proportional controller’s tendency to correct a constant error signal with a derivative controller’s tendency to anticipate changes in the error signal. It achieves this by driving a constant-bearing approach to the target, which typically leads to interception, rather than tail-chasing, of moving targets at *k*_*N*_ *>* 1. This guidance gain *k*_*N*_ is usually called the navigation constant and is conventionally denoted *N*, but we write it here as *k*_*N*_ for consistency with our notation for *k*_*P*_ and *k*_*D*_. In contrast to proportional pursuit, proportional navigation is considered an optimal guidance strategy, in the sense that it minimizes the squared steering effort against non-manoeuvring targets, given appropriate tuning of the navigation constant (27). These properties make proportional navigation an appealing candidate for gap-oriented steering, but previous research on birds has focused only on pursuit-based guidance laws involving the proportional-derivative guidance terms *k*_*P*_ and *k*_*D*_ (20). In fact, proportional navigation can also be used to model deviated pursuit when *k*_*N*_ = 1, which matches the subject’s turn rate 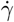 to its line-of-sight rate 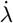, and therefore tends to keep the deviation angle *δ* constant at its initial value *δ*_0_ = *δ*(0) (see Fig. 1). We make use of this fact to avoid having to fit *δ*_0_ as a second free parameter in this single-input guidance law (Table 1).

Different linear combinations of these *k*_*P*_, *k*_*D*_, and *k*_*N*_ terms have been used to formulate mixed guidance laws of the forms shown in Table 1. For example, mixed *k*_*N*_ *k*_*P*_ -guidance has been used to explain the attack trajectories of hawks pursuing manoeuvring targets (8), whereas proportional-derivative *k*_*P*_ *k*_*D*_ -guidance has been used to model pigeons steering through clutter (20). In some cases, additional guidance terms have been proposed that are not based on any underlying guidance or control theory. For instance, the *k*_*P*_ *k*_*D*_ controller used to model pigeons has been tested with the addition of a stabilizing “inertia” term 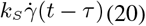. The physical meaning of this term is unclear, however, as using 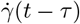 to predict 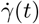 becomes circular as *τ* → 0. For this reason, we do not consider it further here.

Whereas guidance laws can be used straightforwardly to model the target chasing behaviours of birds (6; 8; 7), using them to model clutter negotiation behaviour poses several new challenges. A key contribution of the pioneering work by Lin *et al*. (2014) was to treat clutter negotiation not as an obstacle avoidance behaviour, but as a gap targeting behaviour, thereby enabling its treatment as a classical guidance problem (20). Framing an animal’s movement through a field of obstacles as a sequence of consecutive gap-aiming events (20) avoids the need to model more complex attractor-repeller type behaviours, but risks over-fitting if there are many obstacles, and hence multiple gaps, present. Moreover, a key challenge in applying this framework is the need to define what constitutes a target, and—following from this—the extent to which obstacle avoidance behaviour can be described as target oriented movement at all. This is because an animal’s perception of what constitutes a gap may be different to how an experimenter defines it (**?**). How an animal perceives a gap will depend not only on the visual angles subtended by the objects in the environment (20), but also on background brightness (**?**), and the subjective distance at which the animal treats an object as an obstacle relevant to gap choice. It follows that the gaps between obstacles are virtual constructs that are liable to change as an animal moves.

In this paper, we therefore use a simplified experimental setup to identify the mechanisms of gap choice and gap steering independently. Using a forced binary choice protocol in which one of the gaps was partially obstructed allowed us to test whether the birds flew towards the gap providing the greatest clearance, or whether they used some alternative method of gap selection. Forcing the birds to manoeuvre towards one of two divergent positions also gave us sufficiently varied trajectory data to test alternative guidance laws modelling their steering (Table 1), and to test whether pigeons target the physical centre of the gap between obstacles as Lin *et al*. proposed (20).

## METHODS

### Subjects and training

We used twelve homing pigeons *(Columba livia domestica)* aged from 2 to 10 years as subjects (Supplementary Data S1 and S2). The birds were reared at the John Krebs Field Station, Wytham, Oxford, UK, as members of a free-ranging population provided with *ad libitum* access to food and water in the home loft to which they returned voluntarily. The most experienced non-retired individuals of the population were selected for experiments in order to minimise distress to the animals. Experiments took place over a four week period from mid-November to mid-December 2018. The birds were caught from their loft immediately before the experiment, and released by an experimenter in a large indoor flight hall with one of its ends open to the outside. The birds usually returned directly to their home loft, which was located within 80 m of the exit. Each subject was released in the empty flight hall on five consecutive days prior to experimentation, to ensure familiarity with the location and its relationship to the home loft before introducing any obstructions. The birds’ flight trajectories were only included in the analysis if the bird flew directly through one of the two gaps without landing or loitering (see below), which left us with a sample size of *N* = 10 birds (see Supplementary Data S1 and S2 for exclusions). All testing was approved by the Animal Welfare and Ethical Review Board of the University of Oxford’s Department of Zoology (permit number APA/1/5/ZOO/NASPA), and we monitored the feather condition and flight behaviour of the birds throughout the study for any signs of stress.

### Experimental set-up and protocol

Experiments were undertaken in a flight hall measuring 20.2 m by 6.1 m, with a minimum ceiling height of 3.8 m. The walls of the hall were covered with camouflage netting to provide homogeneous visual contrast. Flicker-free LED lights provided a mixture of direct 4000 K illumination and indirect 5000 K illumination at approximately 1000 lux, which was designed to mimic overcast morning or evening light conditions. Most of the front wall of the building comprised an open roller shutter door providing natural daylight illumination, presenting a much brighter scene than the interior of the flight hall. The ambient weather conditions and position of the sun’s disc were a source of uncontrolled variation in brightness during the experiments. To minimise such variation, experiments were only undertaken during overcast days.

Pigeons were released at 1.2 m height approximately 2.0 m ahead of the back wall, within ±1 m of the midline. From there, they flew through one of two floor-to-ceiling gaps created to either side of a heavy black curtain hung across the hall, 7.0 m ahead of the back wall (Fig. 2A). These symmetric vertical gaps were approximately 1.2 m wide, and hence up to roughly twice the wingspan of the birds. The aperture of the roller door was not visible to the birds at the point of release, but flying through either gap enabled the pigeons to see and reach this aperture, from where they could exit the building (Fig. 2C). Their home loft was located a short distance (approximately 80 m) behind and to the left of the building (Fig. 2B), so was occluded from view until the birds had exited the flight hall.

**Fig. 2.**
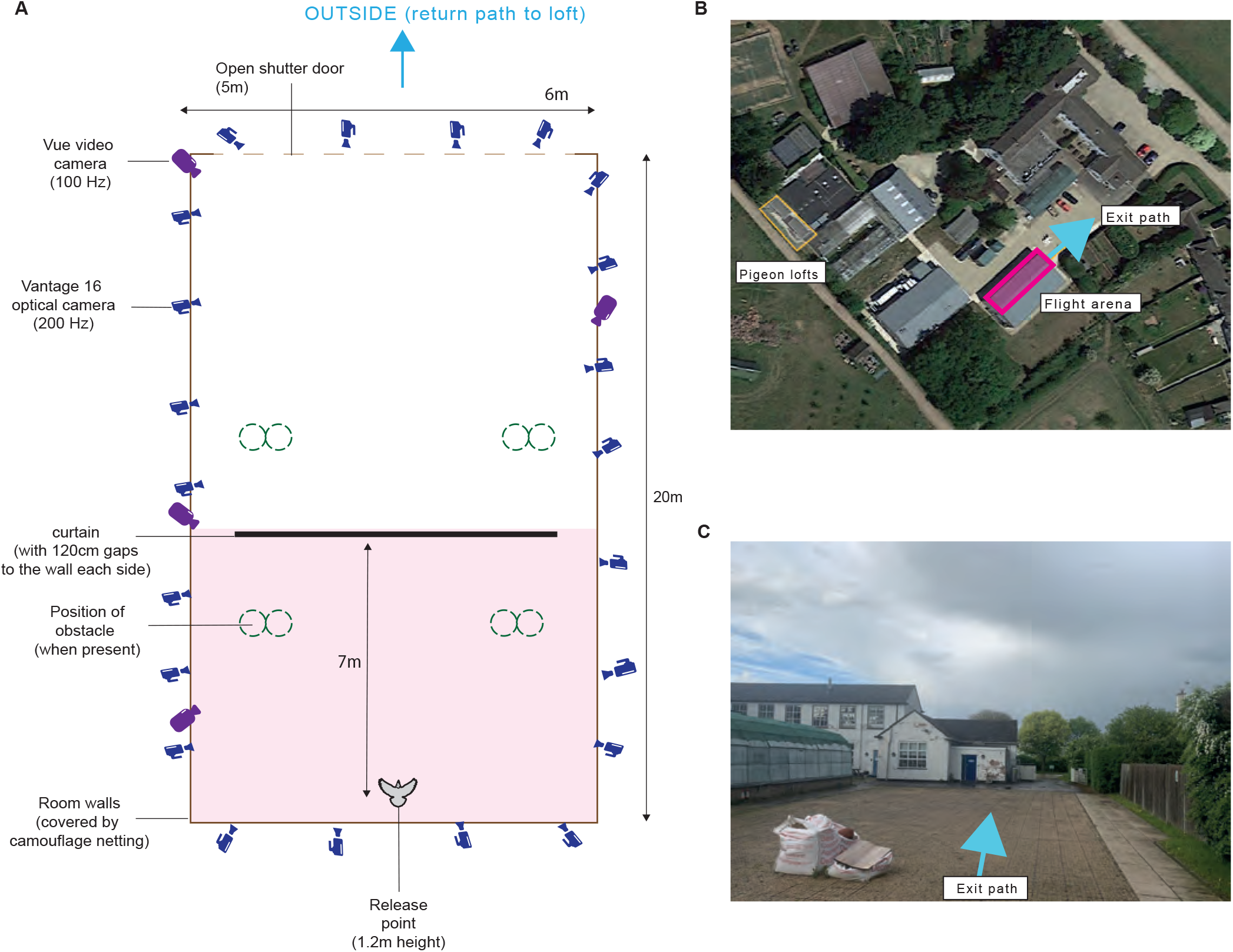
Experimental setup. (A) Flight arena viewed from above. Pigeons flew from the release point to the outside. The path was obstructed by a curtain, creating gaps either side for the pigeons to fly through. The area shaded pink covers the region of interest in which target-oriented steering flight was modelled, from release point to gap. (B) Aerial view of experimental area relative to pigeon home lofts. (C) View out of experimental area from open shutter door. Note that buildings are visible to the left.

We placed a 4.0 m high obstacle in one of two fixed positions in each of the front and back sections of the hall (Fig. 2A). Each obstacle consisted of a pair of 0.3 m diameter white expanded polystyrene cylinders taped together side-by-side, giving high visual contrast against the background. The two obstacles were placed to obstruct the most direct path from (i) the release point to either the right or left gap (back obstacle); and (ii) either the right or left gap of the curtain to the midpoint of the open shutter doors (front obstacle). The second obstacle was not visible to the pigeon until it had reached the gaps on either side of the curtain. We used a randomising procedure to determine where the front and back obstacles were placed on each trial. Here we focus exclusively on the behaviour of the birds in the back section of the hall, between the release point and the curtain. Given that the curtain was located 13.0 m into the hall, we assume that both gaps would have appeared similarly bright to the birds were it not for the placement of the back obstacle (Fig. 2).

On each day of experiments, the pigeons were collected from their home loft, and were fitted with a rigid plate attached to their head using eyelash glue (Duo Quick Set Strip Lash Adhesive Clear Tone). This was fitted just before the start of each recording session, and was removed immediately afterwards. Each plate had a unique asymmetrical configuration of three or four 4 mm diameter spherical retro-reflective markers for motion capture purposes (Vicon Motion Systems Ltd, Oxford, UK). Each pigeon was taken out of its carrier box individually in the flight hall, held for a minute at the release point to acclimatise, and then released to fly freely by the experimenter opening their hands. The next pigeon was only taken out of its carrier box once the preceding bird had exited the flight arena and was out of sight. After all the pigeons had been released, they were collected from the loft to be released again. On a typical test day, we would carry out three releases per pigeon. Testing was only conducted on clear days, and the markers were removed from the pigeons at the end of each day.

### Motion capture

We used an array of 22 high-speed motion capture cameras (Vantage 16, Vicon Motion Systems Ltd, Oxford, UK) and 4 reference video cameras (Vue, Vicon Motion Systems Ltd, Oxford, UK) to record the flights. Cameras were mounted 3.0 m above the floor on scaffolding fixed around the perimeter of the room. The cameras were arranged so that any marker within the recording volume would be visible to at least three cameras, which enabled automatic reconstruction of its 3D coordinates by the motion capture system. Sensor resolution for the Vantage cameras was 4096 × 4096 pixels at a 200 Hz frame rate and 1 ms shutter speed. A strobe unit on each camera emitted infrared light at 850 nm, which falls outside the visible spectrum of the birds (26); an optical filter blocked light at other wavelengths. For the Vue cameras, the sensor resolution was 1920 × 1080 pixels at a frame rate of 100 Hz and a shutter speed of 2 ms.

The camera system was calibrated at the start of each day of experiments. Experiments only went ahead after each camera had been calibrated to an image error accuracy of ≤0.4 pixels for the motion capture cameras, and ≤1.2 pixels for the reference video cameras. Flights were recorded using Vicon Nexus software, which was manually triggered for each pigeon release, with calibration error being checked at the start of each experimental session. The accuracy of the calibration decreased gradually through the day, but the mean three-dimensional reconstruction accuracy for all trials was 0.7 mm (Q1, Q3: 0.57, 1.26 mm). In addition to the retro-reflective markers fitted to the pigeons, we placed 6 mm diameter markers at fixed positions on the edges of the curtain, walls, and obstacles, to record the positions of these features accurately for each trial. These markers were removed temporarily for calibration of the motion capture system.

The Vicon Nexus software outputs the 3D coordinates of all of the markers found within the imaging volume throughout the trial. We labelled each of the recorded markers using custom-written software in MATLAB v9.6 (MathWorks, Inc., Natick, MA, USA), which used clustering algorithms to group markers within and between frames (e.g. to distinguish markers on the pigeon from markers on the obstacles, on the basis of their speeds). The marker labelling procedure was designed to eliminate the false positives that can arise from spurious reflections, but inevitably resulted in some data drop-out; particularly in the less-well visualised part of the flight volume near the ceiling. Marker positions were defined relative to the principal axes of the room, as determined through the calibration of the motion capture system. Any minor discrepancies in the alignment of this global axis system between trials were corrected by applying a Procrustes transformation to the camera coordinates prior to further analysis, so that the global axis definitions for all trials were the same.

### Flight trajectory analysis

We recorded a total of *n* = 105 flights, of which *n* = 96 were judged suitable for further analysis in the sense that the pigeon was recorded flying through one of the gaps without landing or loitering. Of these, *n* = 76 flights were tracked for ≥ 1 s and were used for the purposes of quantitative trajectory analysis. We determined pigeon head position as the mean position of the identified head markers, and fitted a quintic smoothing spline to remove the noise associated with occasional missing or merged markers. The smoothing tolerance was set such that the mean distance between the spline and the data was kept below the maximum span of the head markers, in order to preserve the detail of the head motion. We dropped any flights for which the recorded markers were insufficient for coordinate reconstruction, or where the bird did not fly to the exit point (e.g. perching on a camera instead of exiting the flight hall). Although the trajectory data are fully three-dimensional, we analyse only their horizontal components here, on the basis that the gaps we treat as the targets of the bird’s guidance are extended objects in the vertical dimension.

### Steering controller simulations

We modelled the horizontal steering behaviour of our pigeons using the same trajectory simulation approach used to study target-oriented attack flights in hawks and falcons (6; 8; 7), but tested a broader set of candidate steering controllers comprising all single-input and two-input guidance laws involving linear combinations of *k*_*N*_, *k*_*P*_, and *k*_*D*_ (Table 1). We fitted these guidance laws under two alternative definitions of the target of this gap-aiming behaviour: (i) the midpoint between the curtain edge and the wall (i.e. the point corresponding to the physical centre of the gap); and (ii) the point approximately half a wingspan (0.35 m) into the gap from the edge of the curtain (i.e. the point minimising the clearance with the wings fully spread). The former target definition maximizes the bird’s minimum clearance from the physical structures in its environment, whereas the latter minimizes the clearance on the inside of the turn without risking a collision.

Given the initial conditions recorded for each flight, we simulated the best-fitting two-dimensional flight trajectory commanded under the various guidance models shown in Table 1. For each steering controller, we estimated the guidance parameters and delay for a given flight, or collection of flights, by minimising the mean squared distance between the measured and simulated trajectories over all of the fitted sample points, which we report as the root mean square (RMS) error (*ε*). Whereas we used the guidance law to simulate the bird’s turning, we matched the simulated flight speed to the bird’s measured flight speed, to ensure proper determination of the time derivatives of the line-of-sight angle *λ* and deviation angle *δ*. We did not model deviated pursuit explicitly, because proportional navigation with *k*_*N*_ = 1 simulates a deviated pursuit trajectory in which the deviation angle *δ* remains constant at its initial value *δ*_0_ = *δ*(0), thereby avoiding the need to treat the commanded deviation angle as another free parameter. To account for sensorimotor delay, we lagged the input variables by 0 ≤ *τ* ≤ 0.4 s. The upper end of this range of values exceeds the latency of the pectoralis muscles to the firing of the looming-sensitive neurons in the pigeon’s tectofugal pathway by at least a factor of 4 (30), but is intended to accommodate the possibility that pigeon steering responses in flight might also be delayed by one wingbeat period (approximately 0.15 s inside the flight hall). Taking time *t* = 0 as the first data point, we therefore modelled the bird’s flight trajectory beginning from time *t* = *τ*_*max*_, to ensure that the same section of flight was simulated for all tested values of *τ*. We were thereby able to model each two-dimensional flight trajectory given knowledge of the pigeon’s initial track angle and position, the time history of the pigeon’s speed, and the location of the gap. See Supplementary Information for methodological details and supporting code.

### Statistical analysis

We use hat notation to refer to the least squares estimates of the guidance model parameters (e.g. 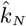 denotes a unique estimate of *k*_*N*_), and tilde notation to denote their median across flights (e.g. 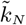 denotes the median value of 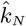 over all flights). We report these medians together with their associated 1st and 3^rd^ quartiles (Q1, Q3), and report the means of quantities estimated directly from the trajectory data together with their standard deviation (mean ± s.d.). Unless stated otherwise, we report two-tailed *p*-values throughout. We use a generalized linear mixed effects (GLME) model with binomial link function in R (25) to analyse the factors affecting gap choice. We use sign tests to test whether the fitted guidance constants were consistently positive or negative, or lower than some critical numerical value (see Discussion). We use Wilcoxon signed rank tests to test for differences in the median RMS error of the fitted guidance models, and use two-sample *t*-tests to test for differences in the means of quantities estimated directly from the trajectory data. We treat each flight as an independent datapoint when analysing the detailed properties of the trajectories, which means that some of the analyses risk pseudo-replication within birds.

## RESULTS

Upon release, the pigeons flew in the direction of the open roller shutter door, even though its aperture was not directly visible from the release point, and despite this being in a different direction to the home loft. Most birds flew directly to the outside via one of the two gaps, but a few began circling in the section of the hall before the curtain. For the purposes of our analysis of gap selection and steering, we only consider data collected between the release point and the curtain (Figs. 2, 3). The mean speed of flights through the unobstructed gap (4.1 ± 0.7 m s^-1^) was significantly higher than for flights through the obstructed gap (3.5 ± 0.8 m s^-1^; two-sample *t*-test: *t*_(50)_ = 2.63, *p* = 0.011). These speeds are slow compared to the cruising speeds of pigeons flying in the open, which can exceed 10 m s^-1^ (24), but are similar to the flight speeds observed previously under similarly cluttered experimental conditions (20).

**Fig. 3.**
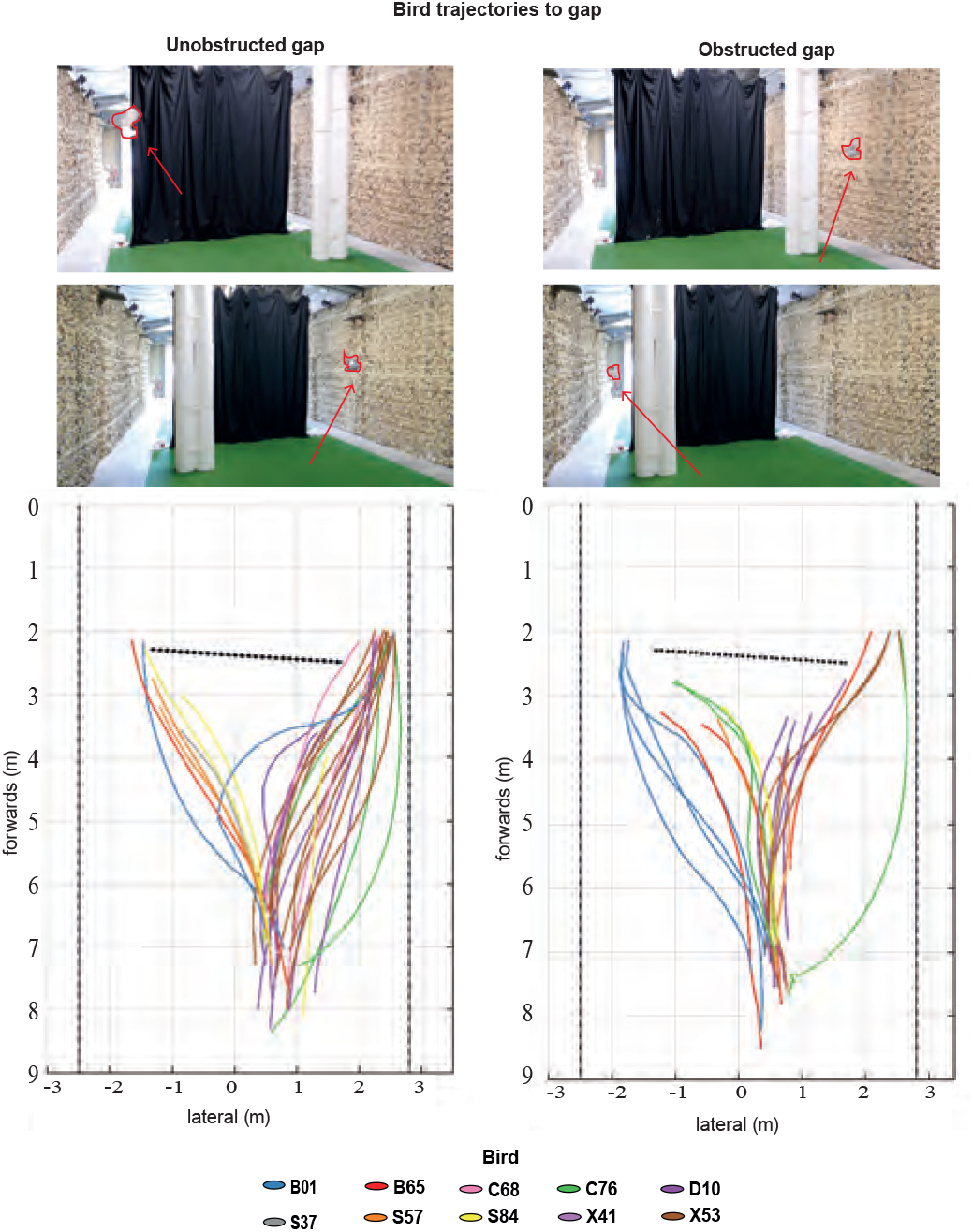
Gap negotiation behaviour in pigeons. All *n* = 96 flight trajectories, colour-coded by individual and viewed from above. Obstacle position was randomised throughout, and pigeons were free to fly through either gap. Trials in which the pigeon flew through the non-obstructed gap (left) are shown separately to trials in which the pigeon flew through obstructed gap (right). Images above show typical flight behaviour and illustrate the experimental setup.

### Pigeons select gaps on the basis of handedness not size

The pigeons usually flew forwards upon release, before steering towards one of the two available gaps (Fig. 3). We interpret this initial flight behaviour as an escape response, with the first unambiguous evidence of gap selection only coming later in the flight. The pigeons flew through the unobstructed gap on 53% of the 96 successful flights, and through the obstructed gap on the remaining 47% (odds of selecting the unobstructed gap: 1.13). The presence of an obstacle had no statistically significant effect on gap choice (GLME: *z* = −1.6, *p* = 0.1), so there is no evidence that the pigeons headed for the unobstructed, and hence larger, of the two gaps. Conversely, there was a clear, statistically significant trend for the pigeons to fly through the right-hand gap (GLME: *z* = 2.1, *p* = 0.03), which they took on 67 out of the 96 flights (marginal odds estimate of selecting right-hand gap: 3.04). Since the home loft was located to the left of the exit, it follows that the birds were not choosing the gap that was most closely aligned with their intended flight direction, but appear instead to be flying with a specific handedness. Importantly, the direction of this handedness was not universal to all birds: some individuals displayed a strong preference for flying towards the right-hand side independent of obstacle placement (e.g. pigeons B01, B65, S57, X41), as indicated by odds ratios significant larger than 1, whereas a smaller number of individuals (e.g. pigeons D10, X53, C68) displayed a strong preference for flying to the left independent of obstacle placement, with odds ratios significantly below 1 (Fig. 4).

**Fig. 4.**
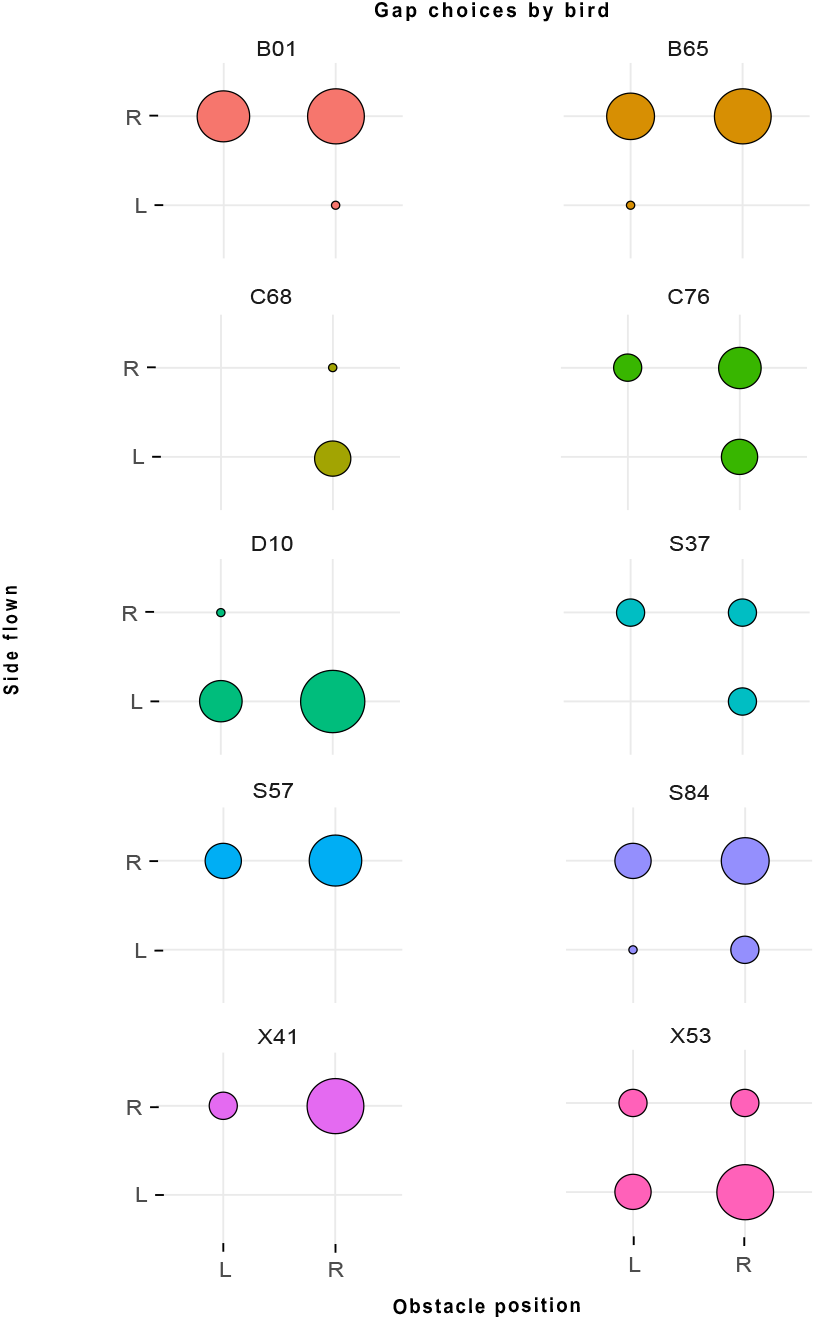
Gap selection behaviour in pigeons. Distribution of gap choice comparing side flown to side on which obstacle was placed for all *n* = 96 flights; area of plotted circle is shown proportional to frequency of choice. Most pigeons display a clear preference for flying on one or other side of the curtain—most often the right-hand side. This idiosyncratic preference does not seem to be affected by the placement of an obstacle between the release point and the gap.

### Simulations of steering behaviour

Because the presence of an obstacle is expected to have modified the bird’s perception of the obstructed gap, we only consider flights taken through the unobstructed gap for the purposes of modelling the steering controller. Because the birds did not necessarily begin their target-oriented steering behaviour immediately upon release (Fig. 3), we only use the data from the last 1.0 s of flight recorded before the pigeon entered the gap for the purposes of model selection and statistical inference, although we extend the fitted section of flight to cover the entire recorded trajectory later. This 1.0 s period of flight does not necessarily extend to the point at which the bird finally reached the gap, owing to marker occlusion in the vicinity of the curtain. With these restrictions, we analysed the subset of *n* = 23 flights from *N* = 10 individuals with ≥ 1 s of continuous recording in which the pigeons were tracked to within ≤ 1 m of the unobstructed gap through which they then flew.

The birds tended to fly upwards from the release point, with a mean altitude gain of 0.73 ± 0.48 m. However, as the task involved flying through vertical gaps that extended from floor to ceiling, we only analyse the birds’ steering with respect to the horizontal. The mean transverse clearance of the pigeons from the curtain was 0.47 ± 0.30 m, or approximately two thirds of an average wingspan, with 0.80 ± 0.48 m mean transverse clearance from the wall. Although the pigeons therefore tended to fly closer to the curtain than to the wall, the fit of the guidance simulations with parameters fitted independently to each flight was closer when the target was defined as the midpoint of the gap than when the target was defined as the point 0.35 m from the edge of the curtain (Wilcoxon signed rank test: *z* = −2.17, *p* = 0.030; test of mean RMS error over all six guidance models fitted independently to each flight). The quantitative results reported below are therefore for simulations targeting the midpoint of the gap, which is the same definition used in previous work on pigeon gap-steering behaviour (20). The quantitative results of the simulations targeting the point 0.35 m from the edge of the curtain are provided as Supplementary Material, and are qualitatively similar in the patterns that they display.

#### Guidance parameters fitted independently to each flight

All six steering controllers that we tested (Table 1) were capable of simulating the flight trajectories closely if their parameters were fitted independently to the last 1.0 s of each flight (Fig. 5A). The associated estimates of the delay spanned a broad range of values (Fig. 6), with a model-average of 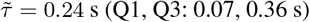 over all six guidance laws. Among the three single-input guidance laws that we tested, *k*_*N*_ -guidance modelled the data most closely (Fig. 5A), with a median RMS error of 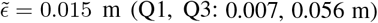. The other single-input controllers that we tested had a higher median RMS error, with 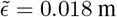 for *k*_*P*_ -guidance (Q1, Q3: 0.011, 0.042 m) and 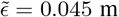 for *k*_*D*_ -guidance (Q1, Q3: 0.020, 0.101 m), but in neither case was this difference statistically significant across flights (Wilcoxon signed rank test: *z* = 0.37, *p* = 0.72; *z* = 1.86, *p* = 0.06). As expected, our estimates of the guidance constants were consistently positive under *k*_*N*_ -guidance (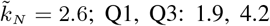; sign test: *p <* 0.001; Fig. 6B), and consistently negative under *k*_*P*_ -guidance (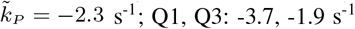; sign test: *p <* 0.001; Fig. 6D). In contrast, they were inconsistently signed under *k*_*D*_ -guidance (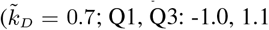; sign test: *p* = 0.40; Fig. 6F), which reflects the over-fitting expected for a controller with no inherent tendency to steer towards a target. In summary, either proportional navigation (i.e. *k*_*N*_ -guidance) or proportional pursuit (i.e. *k*_*P*_ -guidance) is capable of providing a reasonable model of the data, with proportional navigation providing marginally the better fit.

**Fig. 5.**
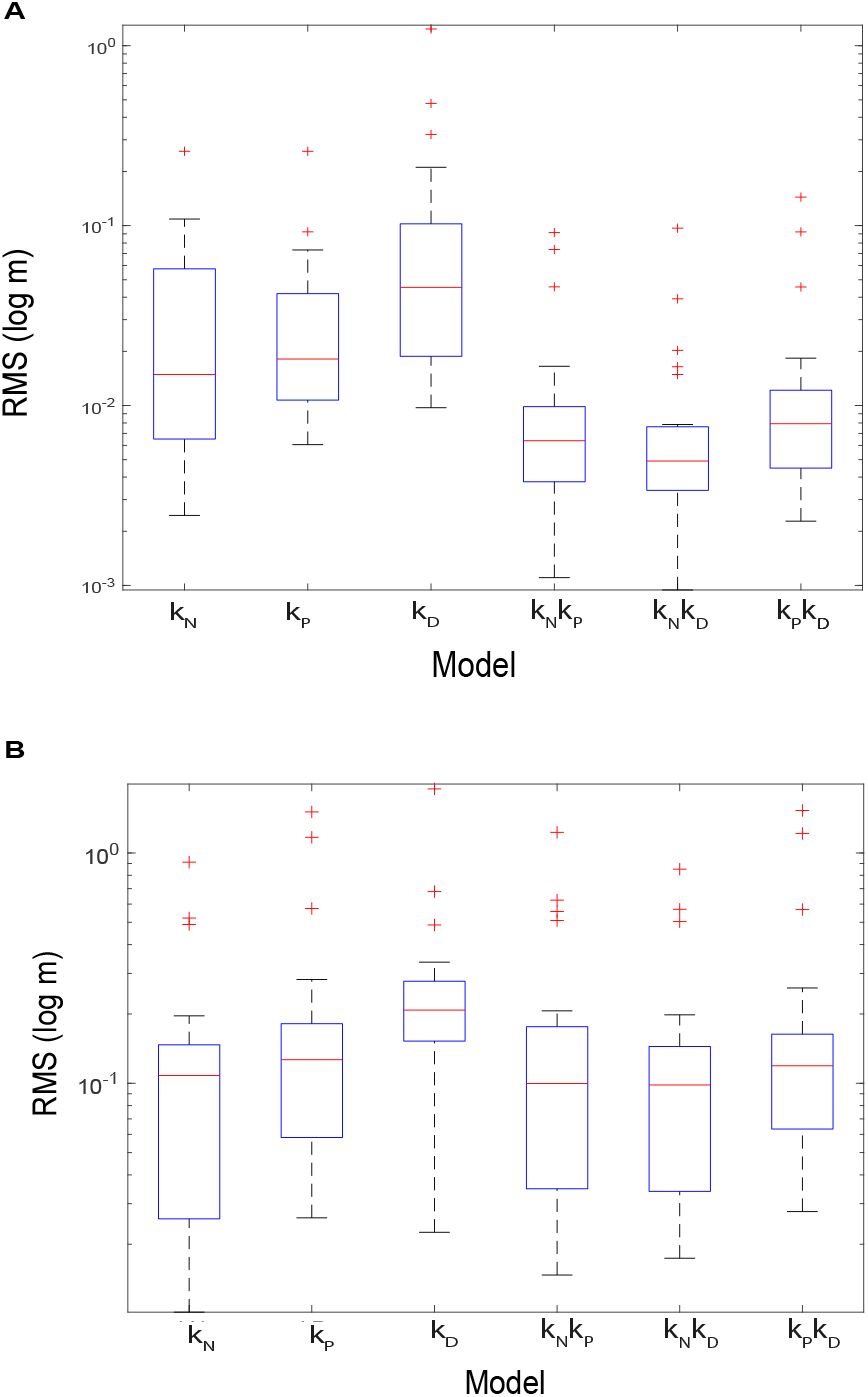
Box plots showing root mean square (RMS) error (*ϵ*) of simulations under each fitted guidance model. (A) Models with parameters fitted independently to each of the *n* = 23 flights through the unobstructed gap. (B) Models with parameters fitted globally to these same flights. The blue box encloses the middle half of the data, with Q1 at the bottom and Q3 at the top; the red line denotes the median; red asterisks denote outliers.

**Fig. 6.**
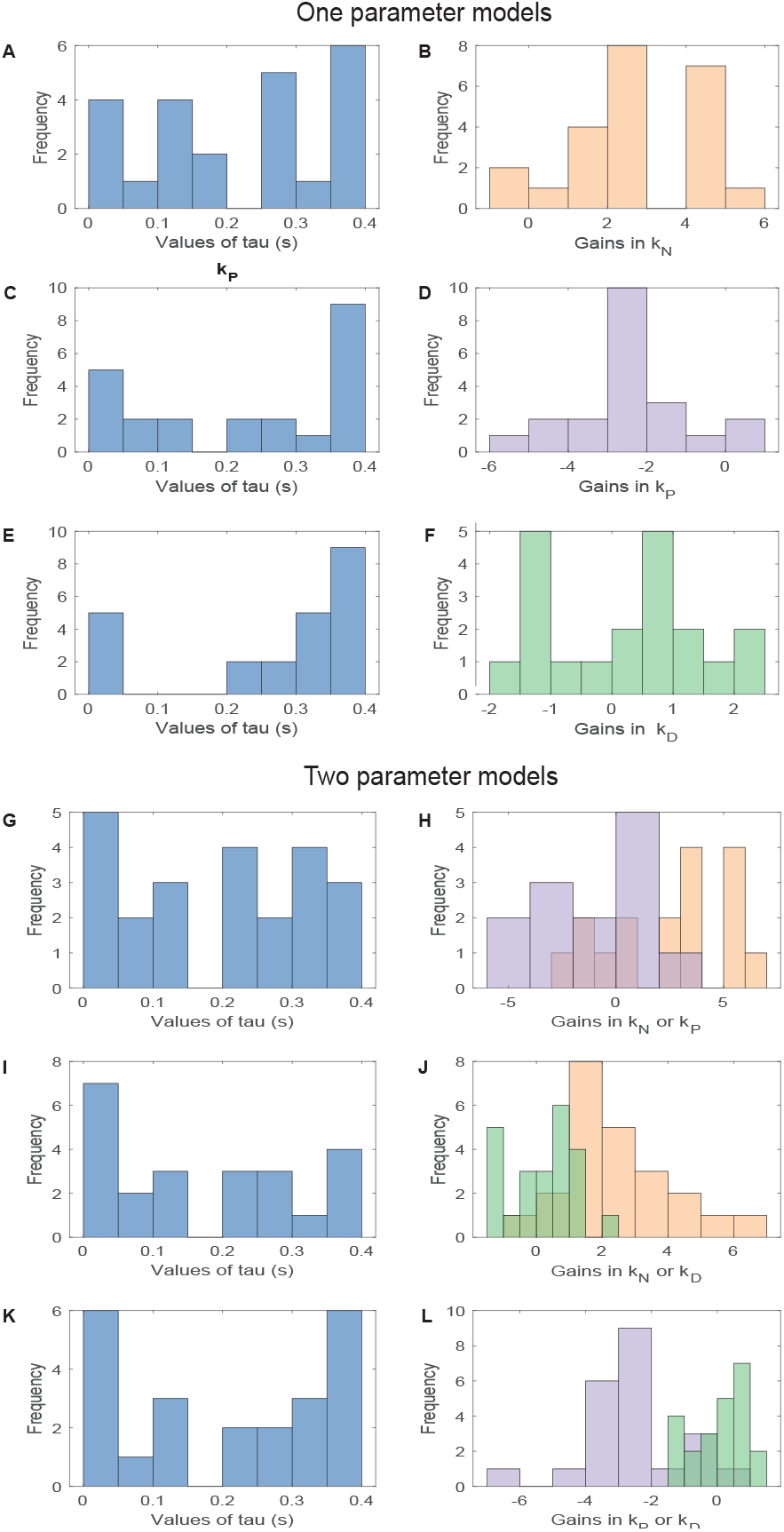
Distributions of parameter estimates for guidance models fitted independently to each flight. Results shown for all *n* = 23 flights through the unobstructed gap, but with some outlying values of the guidance parameters not shown for the few cases where the fitting did not converge on a value within the axis limits (see Supplementary Data S1 for all values including these outliers). Left column: histograms of parameter estimates for time delay *τ* (blue); right column: histograms of parameter estimates for guidance constants *k*_*N*_ (orange), *k*_*P*_ (lilac), and *k*_*D*_ (green), for all *n* = 23 flights. (A,B) *k*_*N*_ -guidance; (C,D) *k*_*P*_ -guidance; (E,F) *k*_*D*_-guidance; (G,H) *k*_*N*_ *k*_*P*_ -guidance; (I,J) *k*_*N*_ *k*_*D*_-guidance; (K,L) *k*_*P*_ *k*_*D*_-guidance.

Introducing a second input variable reduced the error between the measured and simulated trajectories by at least a factor of two (Fig. 5A). Of the various two-input controllers that we tested, *k*_*N*_ *k*_*D*_ -guidance provided the closest fit (Fig. 5A), with a median error of 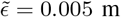 (Q1, Q3: 0.003, 0.007 m) that was significantly lower than for the best-fitting of the single-input guidance laws (Wilcoxon signed rank test: *z* = −4.20, *p <* 0.001). The other two-input controllers that we tested had a higher median RMS error, with 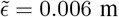 for *k*_*N*_ *k*_*P*_ (Q1, Q3: 0.004, 0.010 m) and 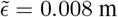 for *k*_*P*_ *k*_*D*_ (Q1, Q3: 0.005, 0.011 m). However, in neither case was this difference statistically significant across flights (Wilcoxon signed rank tests: *z* = 1.00, *p* = 0.32; *z* = 1.43, *p* = 0.15), and in both cases the fit was significantly closer than for the best-fitting of the single-input guidance laws (Wilcoxon signed rank test: *z* = −3.56, *p <* 0.001; *z* = −2.43, *p* = 0.015). Notwithstanding their improved fit over single-input *k*_*N*_ - or *k*_*P*_ -guidance, there is clear evidence that all of the two-input guidance models were over-parameterized. Specifically, the estimates of both guidance constants under *k*_*N*_ *k*_*P*_ -guidance were inconsistently signed (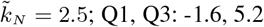; sign test: *p* = 0.21; 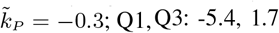; sign test: *p* = 1.0; Fig. 6H). There is therefore no evidence that proportional navigation and proportional pursuit are combined under *k*_*N*_ *k*_*P*_ -guidance. In contrast, although the estimates of *k*_*N*_ and *k*_*P*_ remained consistently signed when fitted in combination with *k*_*D*_, the associated estimates of *k*_*D*_ were inconsistently signed (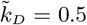 under *k*_*N*_ *k*_*D*_ ; Q1, Q3: -0.6, 1.0; sign test: *p* = 0.40; Fig. 6J; 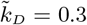 under *k*_*P*_ *k*_*D*_ ; Q1, Q3: -0.5, 0.7; sign test: *p* = 0.40; Fig. 6L). There is therefore no evidence that derivative control is combined with proportional navigation or proportional pursuit under *k*_*N*_ *k*_*D*_ - or *k*_*P*_ *k*_*D*_ -guidance.

In summary, when the guidance parameters were fitted independently to the last 1.0 s of each flight, proportional navigation (i.e. *k*_*N*_ -guidance) gave the best fit of all the single-input steering controllers, but fitted the data only marginally better than proportional pursuit (i.e. *k*_*P*_ -guidance). Whilst it was always possible to achieve a closer fit by adding a second guidance term, the parameter estimates for the second guidance term were always inconsistently signed. The guidance simulations fitted independently to the last 1.0 s of each flight therefore provide no evidence for the involvement of a second guidance term, but do not allow us to distinguish unambiguously between single-input proportional navigation or proportional pursuit.

#### Guidance parameters fitted globally for all flights

Fitting the guidance parameters independently to each flight is appropriate if the true underlying guidance gains vary within or between individuals (6), but risks over-parameterization otherwise. To mitigate this risk, we searched for the unique combinations of parameter settings (Table 2) that minimised the median RMS error for each candidate guidance law over the last 1.0 s of all *n* = 23 flights (Fig 7). We did this using an exhaustive search procedure in which we computed the RMS error on every flight, for all combinations of *k*_*N*_ ∈ [ −0.5, 5.5], *k*_*P*_ ∈ [− 5.5, 0.5], and *k*_*D*_ ∈ [ −2, 2] at 0.1 spacing, and *τ* ∈ [0, 0.4] at 0.005 spacing (SI units). We chose these intervals in light of the ranges of the parameter estimates that we had identified by fitting all flights independently, and used the resulting simulations to identify the global optimum minimising the median RMS error over all flights. Having identified this global optimum at coarse parameter spacing, we then refined the search spacing by a factor of 5 in the vicinity of the coarse global optimum. Fitting the parameters of these steering controllers globally for all flights reduced the total number of fitted parameters by a factor of 23, but also increased the median RMS error by an order of magnitude (Fig. 5B).

**Table 2.** Parameter estimates for steering controllers fitted globally for all flights. Results shown for guidance models fitted to all *n* = 23 flights through the unobstructed gap. Median root mean square (RMS) error 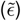 stated with 1st and 3rd quartiles (Q1, Q3) displayed in parentheses. Results for the two-input controllers are stated subject to the constraint that neither guidance constant can be zero.

| model | $\hat{k}_N$ | $\hat{k}_P$ | $\hat{k}_D$ | $\hat{\tau}$ (s) | median RMS error (m) |
| --- | --- | --- | --- | --- | --- |
| $k_N$ | 2.05 | - | - | 0.27 | 0.108 (0.025, 0.146) |
| $k_P$ | - | -1.90 | - | 0.19 | 0.127 (0.058, 0.181) |
| $k_D$ | - | - | -1.00 | 0.01 | 0.208 (0.152, 0.277) |
| $k_N k_P$ | 1.50 | -0.55 | - | 0.29 | 0.100 (0.035, 0.176) |
| $k_N k_D$ | 1.90 | - | 0.10 | 0.24 | 0.098 (0.033, 0.144) |
| $k_P k_D$ | - | -1.90 | -0.10 | 0.27 | 0.119 (0.063, 0.163) |

**Fig. 7.**
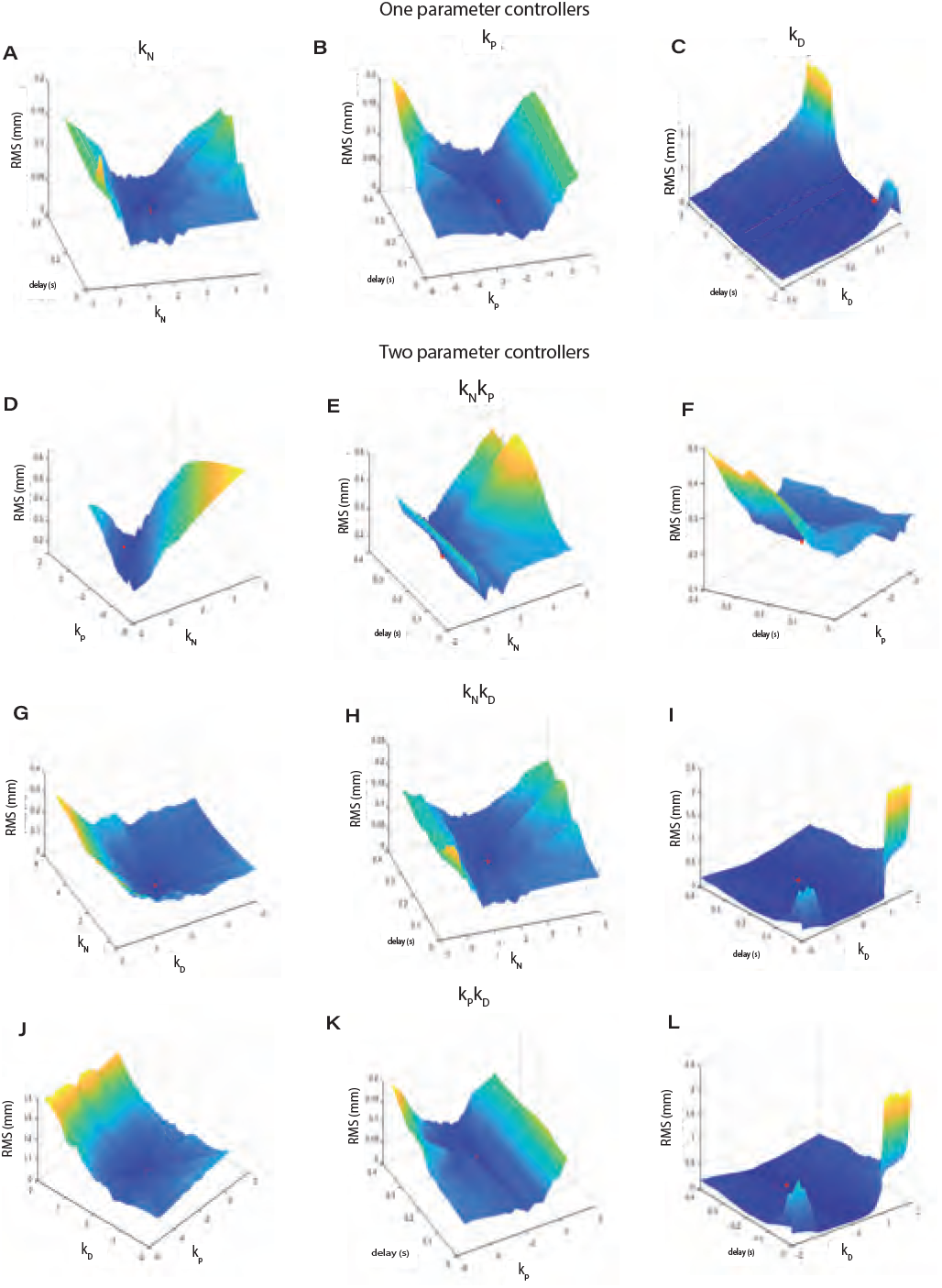
Optimization surfaces for steering controllers fitted globally for all flights. The height of each surface displays the median RMS error for the *n* = 23 flights through the unobstructed gap, as a function of the fitted guidance parameters. (A-C) Single-input guidance laws, plotting median RMS error against the relevant guidance constant (*k*_*N*_, *k*_*P*_, or *k*_*D*_) and time delay *τ*. (D-L) Two-input guidance laws, plotting median RMS error against the relevant pair of guidance constants (*k*_*N*_, *k*_*P*_, and/or *k*_*D*_), or their pairings with the time delay *τ*. Surface colour scaled according to surface height; red dot denotes location of global minimum.

Among the three single-input steering controllers that we tested, *k*_*N*_ -guidance once again provided the best fit to the data, with a median RMS error of 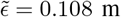 (Q1, Q3: 0.026, 0.144 m; Fig. 5B). The parameter estimate for the navigation constant in this globally-fitted model 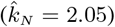 was somewhat lower than the median of the parameter estimates for the navigation constants fitted independently to each flight 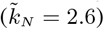, but well within the typical range of fitted values (Q1, Q3: 1.9, 4.2). The associated parameter estimate for the time delay 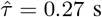 was also in the middle of the range of values fitted independently to each flights (see above). The other single-input controllers that we tested performed significantly less well than *k*_*N*_ -guidance when fitted globally to all fits (Fig. 5B), with a median RMS error of 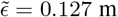 for *k*_*P*_ -guidance (Q1, Q3: 0.058, 0.178 m; Wilcoxon signed rank test: *z* = 2.13, *p* = 0.033) and 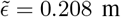 for *k*_*D*_ - guidance (Q1, Q3: 0.156, 0.275 m; Wilcoxon signed rank test: *z* = 2.83 *p* = 0.005).

Adding a second input variable produced only a marginal improvement in the goodness-of-fit of the globally fitted models (Table 2; Fig. 5B). Indeed, although *k*_*N*_ *k*_*D*_ -guidance once again emerged as the best of the two-input models, with a median RMS error of 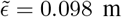 (Q1, Q3: 0.035, 0.140 m), it provided no statistically significant improvement in fit over single-input *k*_*N*_ - guidance in the globally-fitted analysis (Wilcoxon signed rank test: *z* = −0.90, *p* = 0.37). The results of the globally-fitted guidance models therefore confirm without ambiguity that single-input proportional navigation is the best supported of the six alternative steering controllers when fitting these to the last 1.0 s of each flight.

### Extension of model fitting to entire measured trajectory

Having identified delayed proportional navigation targeting the midpoint of the gap as the best supported of the six candidate guidance laws over the last 1.0 s of each flight, we finally used the same model to simulate the entire measured length of the same *n* = 23 trajectories, from time *t* = *τ* to the end of the flight (Fig. 8). Because the values of the navigation constant *k*_*N*_ and time delay *τ* were inherited from the simulations that we had already fitted over the last 1.0 s of each flight (Fig. 6A,B), they are no longer expected to be strictly optimal, and would only be expected to result in a close fit to the data if the birds were engaged in consistent gap-oriented steering behaviour for the entire duration of the measured trajectory. This appears to be the case in general, because with only a few exceptions, the simulations still capture the curvature of the measured flight trajectories closely (Fig. 8), with a median RMS error of 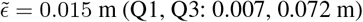.

**Fig. 8.**
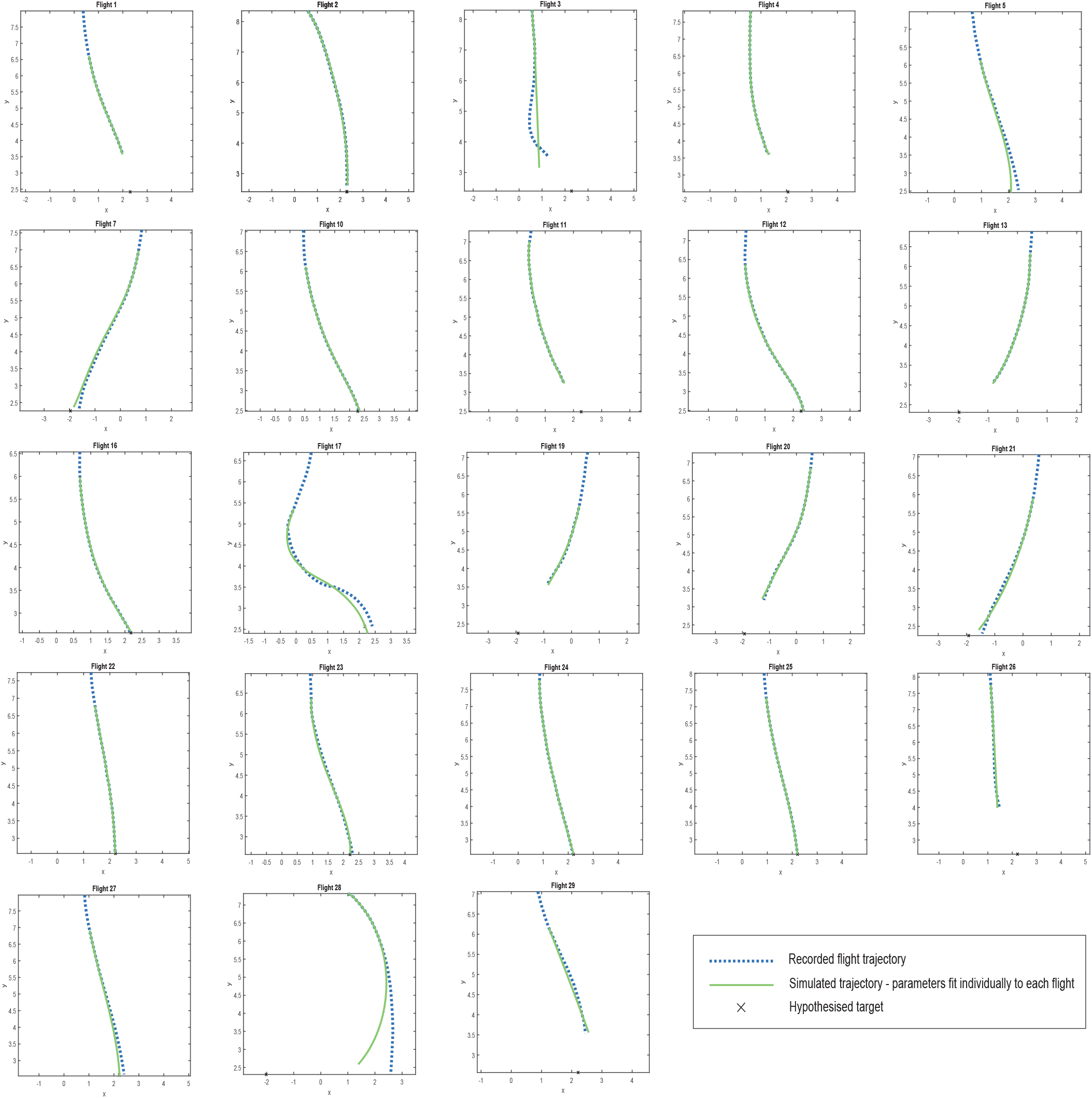
Simulations of gap-oriented steering in pigeons under proportional navigation guidance. Two-dimensional trajectory data (blue dotted lines) are shown for all *n* = 23 flights lasting ≥ 1 s, together with simulations under delayed *k*_*N*_ -guidance (green line) inheriting the values of the navigation constant *k*_*N*_ and time delay *τ* from the simulations fitted independently to the last 1.0 s of each flight. Note that the simulations are plotted beginning from time *t* = *τ* after the start of the recorded trajectory, which is why some trajectories begin with a section of flight for which no simulation is shown.

## DISCUSSION

Our analysis shows that the horizontal flight trajectories of pigeons steering towards vertical gaps are best modelled by the same proportional navigation guidance law that best models falcons attacking stationary or manoeuvring targets (6; 7). There are several lines of evidence that support this conclusion. Firstly, proportional navigation (i.e. *k*_*N*_ -guidance) fitted the trajectory data more closely than any other single-input steering controller that we tested in simulations fitting the guidance parameters independently to each flight, albeit not significantly so at *α* = 0.05. Secondly, proportional navigation fitted the data significantly more closely than any other single-input guidance law that we tested in simulations fitting the guidance parameters globally for all flights. Thirdly, adding a second input to this proportional navigation controller (i.e. adding either a *k*_*P*_ or *k*_*D*_ guidance term) resulted in a marginal and non-significant improvement in the fit when the parameters were fitted globally to all flights, and was associated with inconsistently-signed parameter estimates. Our results further show that the pigeons’ steering behaviour is significantly better modelled by treating the target of its guidance as the midpoint of the gap, rather than the point falling approximately half a wingspan from the nearside edge. This result holds on average across all of the guidance models that we fitted independently to each flight, and also holds for the proportional navigation controller in particular—albeit that the qualitatively better fit of the model treating the midpoint of the gap as the target of the bird’s guidance is not quite significant in this specific case (Wilcoxon signed rank test: *z* = −1.89, *p* = 0.059).

Our results therefore support Lin *et al*.’s conclusion (20) that feeding back the rate of change of the deviation angle (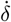) through derivative control (*k*_*D*_) is unimportant to modelling gap steering behaviour in pigeons. However, they also show that this behaviour is better modelled by a proportional navigation (*k*_*N*_) controller feeding back the line-of-sight rate 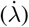 of the midpoint of the gap, than by a proportional pursuit (*k*_*P*_) controller feeding back the deviation angle of the midpoint of the gap (*δ*) as Lin *et al*. originally proposed. This conclusion has implications for our assumptions regarding the underlying sensory mechanism. Specifically, whereas measurement of the deviation angle (*δ*) is expected to require knowledge of the retinal coordinates of the target and angular position of the head with respect to the body, measurement of the line-of-sight rate 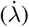 is expected to require knowledge of the retinal drift rate of the target and angular rate of the head in an inertial frame of reference. The sensory requirements of these two guidance laws are therefore quite different, although either could in principle be used to model the data satisfactorily. More importantly, our results confirm the possibility of uniting the study of target-oriented steering behaviours of all kinds under one common algorithmic framework. Specifically, we have now shown that the same proportional navigation guidance law that best models attack behaviours in falcons (6; 7) also provides the best model of gap-oriented steering behaviour in pigeons. This is consistent with the emerging hypothesis that target-oriented guidance behaviours of birds of all kinds share a common evolutionary origin deep in their phylogeny (8).

In fact, our numerical estimates of the navigation constant *k*_*N*_ in the proportional navigation simulations fitted independently to the obstacle avoidance flights of our pigeons (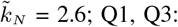 1.9, 4.2) were quantitatively similar to those found previously in peregrine falcons attacking stationary targets 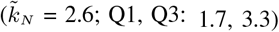. This has several further implications, which we elaborate here by making use of some classical results describing the behaviour of proportional navigation at different values of the navigation constant *k*_*N*_ (27). The statistically significant result that 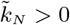 in the guidance models fitted independently to each flight (one-tailed sign test: *p <* 0.001) confirms that the pigeons engaged in target-oriented steering behaviour immediately before passing through the gap, because proportional navigation produces turning towards a stationary target at *k*_*N*_ *>* 0. Likewise, the statistically significant result that 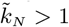 in these simulations (one-tailed sign test: *p <* 0.001) confirms that the pigeons were not using any form of pure or deviated pursuit, which is the flight behaviour that proportional navigation describes at *k*_*N*_ = 1. More generally, proportional navigation produces a turn of ever-decreasing radius against a stationary target at *k*_*N*_ *<* 2, with a turn of constant radius produced at *k*_*N*_ = 2. Hence, the statistically significant result that 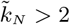 in the guidance models fitted independently to each flight (one-tailed sign test: *p* = 0.047) confirms that the pigeons usually made a turn of ever-increasing radius, thereby straightening out on final approach. This is clearly an appropriate behaviour for flight aimed at negotiating a narrow gap. Proportional navigation at *k*_*N*_ *>* 2 may therefore be a *better* guidance law for negotiating a narrow gap than the previously proposed (20) alternative of proportional pursuit which leads to a tightening turn. Indeed, the median value of 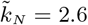 that we found is not significantly lower (sign test: *p* = 0.11) than the theoretical optimum of *k*_*N*_ = 3 that commands the most efficient turn towards a stationary target, in the sense of minimising the control effort measured by the time integral of the squared acceleration command.

It is interesting to note that the goblet-shaped spatial distribution of our pigeons’ flight trajectories (Fig. 3) closely resembles the results of experiments from flies and locusts presented with a choice between two spatially distinct options (28). In both these cases, the animal appears to adopt the average path between the options until some critical phase transition occurs, at which point the system decides spontaneously between them. Our pigeons likewise displayed a centered response initially, flying forwards from the point of release, before turning in the direction of one of the two gaps and then steering through it (Fig. 3). Theoretical modelling of this two-option spatial decision making process (28) associates the moment at which turning begins with the timing of the decision to turn towards one or other option, but the results of our guidance modelling suggest a more nuanced interpretation in the case of our pigeons. Specifically, because the line-of-sight rate of a stationary target will only be non-zero if the subject is moving, it follows that the subject must already be moving in order to generate a steering command under proportional navigation. In the presence of sensorimotor delay, this means that the animal may not begin turning at all until after it has begun moving. Hence, whereas proportional navigation can only produce monotonic turning towards a stationary target in the absence of delay, it can produce a sinuous turn in the presence of a delay. This is already sufficient to explain the sinuous nature of some of the measured trajectories plotted in Fig. 8. Nevertheless, as these trajectories do not begin from the moment of take-off, it remains possible that the pigeons did not make their targeting decision until after they had already begun flying forwards from the release point. Any such delay in their decision making may also explain why the pigeons did not show any clear preference for flying through the unobstructed gap (Fig. 4). This is because the obstacle was placed between the release point and the gap (Fig. 2), and may therefore not have obstructed either gap from the perspective of the subject at the point where its targeting decision was finally taken (Fig. 3).

The preceding account points to the importance of determining both the outcome and the timing of targeting decisions when modeling movement through complex environments. In cluttered environments requiring multiple targeting decisions, the timing of every decision will be critical to determining the overall shape of the trajectory. The general question of how a chain of spatial decisions structures trajectories through complex environments clearly merits further investigation, because the portion of a trajectory over which steering can occur under closed-loop guidance is obviously limited to the interval between the point at which a targeting decision is made and the point at which the animal reaches its target. Fast decision making is therefore critical when flying through clutter, and open-loop turning commands may be important to ensuring that flight is directed appropriately during the period of any delay between target acquisition and the onset of closed-loop guidance. One method of increasing the speed of decision making when choosing between similar alternatives is to introduce a bias in the decision making process. There is clear evidence of biased choice in our data, because at the population level our birds displayed a clear preference for selecting the right hand gap, whilst at the individual level most birds displayed their own idiosyncratic preference for choosing the right or left gap. This is consistent with individual biases observed in budgerigars choosing between two apertures of differing sizes (3). We propose that such biases may aid decision making when flying through clutter, which is a situation in which making and committing to a decision quickly may be at least as important as the decision that is made. This hypothesised enhancement of speed and safety when flying through clutter has been shown through mathematical models to be advantageous for flocks of birds as well as individual (3). How might such biased choices arise? There is widespread evidence of handedness in animals (31), which has already been shown to accelerate homing over longer distances in swimming and flying animals (2). Handedness can be of mechanical, sensory, or neural origin, but based on the information available here, we cannot distinguish whether the biased choice that we observed reflects the intrinsic handedness of different individuals, or an idiosyncratic response to extrinsic cues. For instance, as the preference to fly down the right hand side of the hall continued after the pigeons had passed through the gap, it is plausible that individual birds simply used their visual memory to recapitulate the fine scale route that they had followed on their initial release, as has already been shown in large-scale homing behaviours (4).

Finally, the guidance framework that we have applied has implications for understanding the sensory information that animals use to structure their goal-directed behaviours. The steering behaviour that we observed in our pigeons was best explained by assuming that they targeted the midpoint of the gap (which would maximise the clearance on both sides), rather than the point half-a-wingspan in from the near edge (which would have minimised the nearside clearance with the wings spread). It is therefore reasonable to assume that this behaviour would not have required metric estimation of the physical width of the gap. On the other hand, as the midpoint of the gap represents a virtual target, it is an open question how its line-of-sight rate 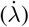 would have been estimated by the birds. One possibility is that the birds could have centred the gap in their visual field, and estimated its line-of-sight rate from the angular rate of their head by integrating the angular accelerations sensed by their vestibular system. Another possibility is that they could have made use of the rotational optic flow cues produced by the head’s self-motion relative to a fixed visual background. A gap-centering response of this kind would differ from the mechanism observed in birds avoiding single obstacles, which appear to fixate their gaze on the edge of the obstacle that they are aiming to avoid (18; 22). This again raises the question of whether animals perceive clutter as a set of gaps to be aimed at, or a set of obstacles to stay clear of. Establishing a guidance model for gap-oriented steering behaviour in pigeons, as we have done here, reduces the problem to one of determining how and when they select which clearances to fly through.

### Limitations

Although the number of individuals that we tested was twice that of other similar studies (3; 20), our sample size is small in absolute terms, and involves repeated measures from the same individuals that could not always be controlled for statistically. It follows that our results cannot necessarily be assumed to generalise more broadly. The sections of flight that were fitted under the guidance models are also quite short, which is inevitable when modelling flight through clutter, but explains why the observed flight trajectories can often be modelled successfully by more than one guidance model. Hence, although proportional navigation was the best supported of the various guidance models that we tested, it is plausible that a different result might emerge given a larger sample size or longer flight duration. Future studies could therefore usefully increase the distance between the release point and the gaps to be traversed, as well as the total number of individuals tested. Finally, it is important to note that the experimenter who released the birds could not be made blind to the experimental condition (Fig. 2), but as there was no evidence of any effect of obstacle placement on gap choice, this is unlikely to have affected the experimental results. Other possible biases such as the handedness of the experimenter would have affected different birds similarly, so are unlikely to explain their idiosyncratic preference for choosing the right or left gap.

## Acknowledgements

We thank Lucy Larkman for her assistance with the bird experiments and animal husbandry. We thank Marco Klein-Heerenbrink and Sofia Minano-Gonzalez for their assistance with data processing.

## Contribution

Conceptualization: N.P.-C., G.T.; Methodology: N.P.-C.; Investigation: N.P.-C.; Software: N.P.-C., G.T.; Data curation: N. P.-C.; Formal Analysis: N.P.-C., G.T.; Visualization: N.P.-C.; Writing - original draft: N.P.-C. Funding acquisition: N.P.-C., G.T.; Project administration: G.T.; Supervision: G.T.; Writing - review & editing: G.T.

## Data availability

Flight trajectory data and code is available on figshare at https://doi.org/10.6084/m9.figshare.19285289.

## Supplementary

Electronic supplementary material is available online and includes the results of the simulations fitted independently to each flight or globally to all flights. **Data S1** contains the results of simulations treating the midpoint of the gap as the target of the bird’s guidance. **Data S2** contains results of simulations treating the point 0.35 m from the edge of the curtain as the target ofthe bird’s guidance.

